# Neural dynamics of reselecting visual and motor contents in working memory after external interference

**DOI:** 10.1101/2024.12.13.628347

**Authors:** Daniela Gresch, Larissa Behnke, Freek van Ede, Anna C. Nobre, Sage E.P. Boettcher

## Abstract

In everyday tasks, we must often shift our focus away from internal representations held in working memory to engage with perceptual events in the external world. Here, we investigated how our internal focus is reestablished following an interrupting task by tracking the reselection of visual representations and their associated action plans in working memory. Specifically, we ask whether reselection occurs for both visual and motor memory attributes and when this reselection occurs. We developed a visual-motor working-memory task in which participants were retrospectively cued to select one of two memory items before being interrupted by a perceptual discrimination task. To determine *what* information was reselected, the memory items had distinct visual and motor attributes. To determine *when* internal representations were reselected, the interrupting task was presented at one of three distinct time points following the retro-cue. We employed electroencephalography time-frequency analyses to track the initial selection and later reselection of visual and motor representations, as operationalized through modulations of posterior alpha (8-12 Hz) activity relative to the memorized item location (visual) and of central beta (13-30 Hz) activity relative to the required response hand (motor). Our results showed that internal visual and motor contents were concurrently reselected immediately after completing the interrupting task, rather than only when internal information was required for memory-guided behavior. Thus, following interruption, we swiftly resume our internal focus in working memory through the simultaneous reselection of memorized visual representations and their associated action plans, thereby restoring internal contents to a ready-to-use state.

**Significance statement:** A key challenge for working memory is to maintain past visual representations and their associated actions while engaging with the external environment. Our cognitive system must, therefore, often juggle multiple tasks within a common time frame. Despite the ubiquity of multi-task situations in everyday life, working memory has predominantly been studied devoid of additional perceptual, attentional, and response demands during the retention interval. Here, we investigate the neural dynamics of returning to internal contents following task-relevant interruptions. Particularly, we identify *which attributes* of internal representations are reselected and *when* this reselection occurs. Our findings demonstrate that both visual and motor contents are reselected immediately and in tandem after completion of an external, interrupting task.

## Introduction

Successful behavior often requires maintaining internal representations while simultaneously handling external sensory events. For example, when cycling to a new ice cream shop, we must retain a memorized set of directions while responding to external events, such as navigating around potholes in our path. While earlier research has predominantly focused on how sensory information is encoded, maintained, and retrieved in working memory (Brady et al., 2011; Luck & Vogel, 2013), recent studies have begun to explore the mechanisms supporting the protection of internal visual representations against external interference (Bettencourt & Xu, 2016; Degutis et al., 2024; Gresch et al., 2021, 2022; Hakim et al., 2020; Hallenbeck et al., 2021; Kreither et al., 2022; Lorenc et al., 2018, 2021; Xu, 2017, 2024; Zickerick, Rösner, et al., 2021). A critical question that remains, however, is how our internal focus is reestablished after engaging with interference – specifically, *what* internal information is reselected and *when* this reselection process occurs.

It is crucial to recognize that working memory in real life is more than a short-term storage for detailed sensory information. Rather, past sensory representations must be considered alongside their prospective actions. Internal representations inherently represent the past, however, the purpose of holding sensory information in memory is to guide future behavior (Heuer et al., 2020; Myers et al., 2017; Nobre & Stokes, 2019; Olivers & Roelfsema, 2020; van Ede, 2020; van Ede & Nobre, 2023). Consistent with this notion, contemporary studies have begun demonstrating that encoding, maintaining, and selecting visual information in working memory often occurs alongside the planning of associated actions (Boettcher et al., 2021; Echeverria-Altuna et al., 2024; Ester & Weese, 2023; Nasrawi et al., 2023; Nasrawi & van Ede, 2022; Schneider, Barth, & Wascher, 2017; van Ede, Chekroud, Stokes, et al., 2019).

To date, limited research has examined motor signatures in scenarios where maintenance is interrupted by external interference. Current evidence suggests that while action plans are reinstated in working memory following task-relevant interruptions (Boettcher et al., 2021), the extent of this reinstatement is reduced compared to situations where no interference occurs (Ülkü et al., 2024a, 2024b; Zickerick, Kobald, et al., 2021). Although this work has helped clarify whether action representations are reselected following interference, important theoretical gaps remain. First, it is unclear whether visual-spatial information is additionally reinstated. This would not be strictly necessary if participants had already extracted the target’s relevant features and exclusively prepared the required motor plan for memory-guided behavior. Second, the temporal dynamics of internal reselection remain unclear. Two possibilities arise: reselection could occur either just in time for the upcoming working-memory task or immediately after the interfering event, regardless of when the memory content is needed for behavior.

If both visual and motor signals are reselected, a third question arises regarding the temporal relationship between these two signatures. Recent work has demonstrated that when memorized visual information is probed to guide behavior following a blank memory delay, action plans can be selected from working memory concurrently with the probed visual representations (van Ede, Chekroud, Stokes, et al., 2019). However, it is not known whether the reinstatement of visual and motor attributes also concurs when endogenously triggered by engagement with external interference.

To address what (just motor or visual and motor) and when (just in time or immediately) internal representations become reselected after engaging in an external task, we recorded electroencephalography (EEG) to track the neural dynamics of visual information and prospective manual actions in a task that systematically manipulated the onset of an interrupting task during the working-memory delay. Contra-versus ipsilateral modulations of posterior alpha (8–12 Hz) and central beta (13–30 Hz) activity served as markers for the reselection of sensory- and action-related contents in working memory, respectively. To preview our findings, we demonstrate that both visual and motor signatures are reselected immediately after responding to the interrupting task. Moreover, visual and motor reselection occurs in tandem.

## Materials and Methods

### Participants

The study was approved by the Central University Research Ethics Committee of the University of Oxford. Thirty-four volunteers participated. Two participants were excluded following an *a priori* behavioral trial-removal procedure (see Behavioral analysis). Furthermore, two datasets were excluded due to the absence of eye-tracking data, which made it impossible to verify that participants kept their gaze fixed during the task (see Eye-tracking analysis). The remaining 30 participants (age range: 20 to 40; mean age: 26.2; gender: 16 female, 14 male; handedness: 25 right-handed, 5 left-handed) had normal or corrected-to-normal vision. Individuals provided written informed consent before participating in the study and were compensated at a rate of £20 per hour.

### Task and procedure

The experimental design was adapted from a recently developed visual-motor task (Boettcher et al., 2021; Echeverria-Altuna et al., 2024; Nasrawi et al., 2023; van Ede, Chekroud, Stokes, et al., 2019). On each trial, participants memorized the orientations of two differently colored bars (**Fig. 1A**). At the end of each trial, participants reproduced the orientation of one of these bars, indicated by a correspondingly colored retro-cue which appeared during a memory delay. A key feature of this task was that each bar’s orientation was directly linked to the response hand required to reproduce its tilt. Since the prospective response hand and the cued item’s location were manipulated orthogonally, the task design could disambiguate EEG markers of motor selection (i.e., required response hand) from those of visual selection (i.e., location of memorized item) following the retro-cue onset.

**Figure 1.**
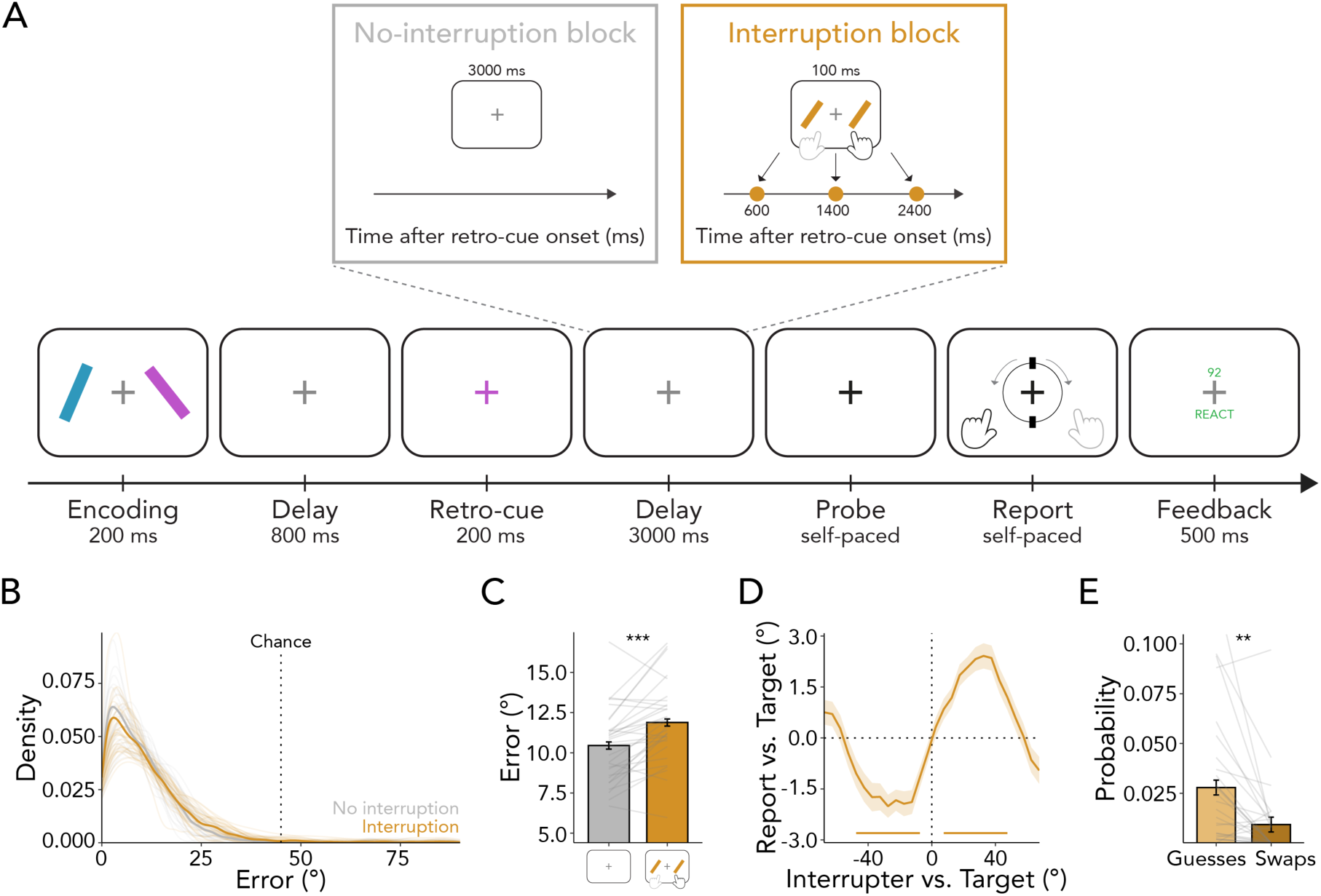
Task schematic and performance. **(A)** Trials started with an encoding display consisting of two lateralized tilted bars. Participants’ task was to remember the angle of both bars, of which one had to be reported at the end of the trial. After a delay, a retro-cue appeared indicating the to-be-reported item. In most blocks, interrupter items occurred for 100 ms during the memory delay following the retro-cue. In this interrupting task, participants had to indicate whether items were tilted to the left or right. The interrupter items occurred at three distinct time points within a block (i.e., at 600 ms, 1400 ms, or 2400 ms after retro-cue onset; at 400 ms, 1200 ms, 2200 ms after delay onset). The delay between retro-cue offset and probe onset was always 3000 ms long. After the delay, the central fixation cross turned black, indicating that the previously retro-cued item should now be reported. Participants reported the memorized tilt using a response dial and were provided feedback at the end of the trial. **(B)** Distribution of reproduction errors in the working-memory task for no-interruption and interruption blocks. Errors in both conditions deviated from chance level. **(C)** Reproduction errors in the working-memory task were smaller for no-interruption (gray bar) compared to interruption blocks (orange bar). **(D)** Average response bias relative to the angular difference between the target and interrupting items revealed a working-memory bias towards the interrupters (i.e., an attractive bias). **(E)** The proportions of guesses and swaps with the interrupting items were low, with swaps occurring less frequently than guesses. Error bars and shadings indicate *M* ± *SEM*. Lines indicate individual participants. * indicates *p* < 0.05, ** indicates *p* < 0.01, *** indicates *p* < 0.001.

To investigate the reselection of internal visual and motor signatures, an external, interrupting task was included in the majority of blocks following the presentation of the retro-cue (block type: no-interruption block vs. interruption block). This external task could occur at one of three potential time points: either 600 ms, 1400 ms, or 2400 ms after the onset of the retro-cue (interruption onset: early vs. medium vs. late). For trials within a fixed block, the external task occurred at only one of these three time points; whereas for trials within a variable block, the external task could occur early, medium, or late with equal likelihood (temporal predictability: fixed block vs. variable block).

At the start of each trial, two tilted bars were simultaneously presented against a gray (RGB value: [102, 102, 102]) background for 200 ms. One bar was always positioned to the left and the other to the right of a central, white fixation cross (RGB value: [255, 255, 255]). Independent of the location, one of the bars was tilted to the left (anticlockwise) and the other to the right (clockwise). To avoid angles being too close to the vertical and horizontal meridians, the items’ angles were randomly drawn in increments of ±1° between ±10° and ±80°. Within each block, a bar oriented leftwards or rightwards was equally likely to appear in either the left or the right position. The stimuli were each approximately 0.7° in width and 5° in length, centered at a viewing distance of 5.5° visual angle left or right from fixation. On each trial, the colors of the memory items were randomly drawn from a set of three highly distinguishable colors (RGB values: blue [0, 178, 255], orange [239, 169, 33], and pink [255, 0, 246]). At encoding, both lateralized items were equally likely to be probed, rendering them equally relevant.

Visual encoding was followed by a memory delay of 800 ms, during which the fixation cross remained on the screen. After this delay, the central fixation cross changed color for 200 ms to match one of the previously presented bars. This retro-cue was 100% valid, always indicating the to-be-reported item.

Retro-cue presentation was followed by a delay interval of 3000 ms, during which the white fixation cross was presented. The delay period could be blank or include an interrupting task, depending on the block. Two-thirds of blocks included an interrupting task: two tilted bars were simultaneously presented for 100 ms, appearing in the same size and locations as those in the memory display. Both bars in this external task had the same orientation and color. Their angle was pseudo-randomly drawn in increments of ±1° between ±10° and ±80°. Their leftwards and rightwards tilts were counterbalanced within each block, and their color always differed from those of the two preceding memory items. If the bars were tilted to the left, participants pressed the F key with their left index finger. If the bars were tilted to the right, participants pressed the J key with their right index finger.

To track *when* internal representations were reselected – immediately or just in time – the interrupting task could occur at three distinct onsets. Across separate blocks, interference was introduced at a fixed onset time, occurring either at an early (600 ms), medium (1400 ms), or late time (2400 ms) relative to the retro-cue onset (i.e., 400 ms, 1200 ms, or 2200 ms after delay onset). Moreover, we included variable onset blocks in which interrupters were equally likely to occur at any of these three time points. The total length of the memory delay was always 3000 ms, regardless of whether and when the external task appeared. In the late condition, the response window for reacting to the interrupter was 800 ms, providing participants sufficient time to report its orientation.

Following the memory delay, the fixation cross changed to black (referred to as the “probe”), indicating that the previously cued item should now be reported. The probe was deliberately presented in a non-target color to ensure participants utilized the retro-cue when performing the task. Participants were never probed about the interrupter items. Following the appearance of the probe, participants had unlimited time to decide on their response. To report a leftwards (rightwards) angle, participants were asked to press the F (J) key on the keyboard using their left (right) index finger. After response initiation, a visual response dial was displayed on the screen, always starting in vertical position. The response dial had the same diameter as the length of the bars (5°) and always appeared surrounding the fixation.

The dial rotated leftwards when the F key was pressed and rightwards when the J key was pressed (either by holding the key down or pressing the key repeatedly) at a speed of 1/10° per millisecond. Critically, the dial could only be rotated in the direction that was initially indicated by the participant. For example, if a participant started pressing the F key after the probe, the dial would only move leftwards, and it would not be possible to move the dial back towards the right with the J key. Since the response dial always started in a vertical position and could not be rotated beyond ±90°, a leftwards (rightwards) oriented bar could only be correctly reported with a left (right) key. Consequentially, the hand required for responding was directly linked to the angle of the bar that was probed. Once the response dial aligned with the memorized tilt of the memory item, participants pressed the space bar to verify their response. Critically, because the tilt and location of the cued item were counterbalanced (i.e., a left- or right-positioned memory bar was equally likely to be paired with a left- or right-hand response in the working-memory task), it was possible to independently characterize neural activity reflecting the target item’s memorized location and the prospective response hand associated with the target item’s tilt. Additionally, we orthogonally manipulated the tilt of the cued memory item and the tilt of interrupter items across trials (i.e., a left- or right-hand response in the working-memory task was equally likely paired with a left- or right-hand response in the external, interrupting task).

After a 100-ms delay in which only the fixation cross stayed on the screen, participants received a 500-ms feedback regarding their working-memory and external-task performance. Working-memory performance was indicated by a number ranging from 0 to 100, with 100 indicating a perfect report and 0 indicating that the adjusted orientation was perpendicular to the angle of the target item. If working-memory performance was below 75, the number was presented in red font (RGB value: [255, 0, 0]), otherwise appearing in green (RGB value: [0, 240, 15]). This feedback was explicitly explained to participants and was used to motivate participants to respond accurately to the working-memory item. When participants took longer than 800 ms to respond to the external task, responded using the wrong key, or did not respond at all, the word “REACT” was displayed in red font, otherwise, the word “REACT” appeared in green font. This motivated participants to respond accurately and quickly to the interrupting task. In no-interruption blocks, the latter feedback was not presented. Trials were separated by an inter-trial-interval randomly drawn between 750 and 1000 ms. Between blocks, participants received additional feedback about their average accuracy in the interrupting and working-memory tasks of the previous block.

The experiment consisted of 648 trials divided across 27 blocks, each including 24 trials. Of the 27 blocks, 9 were no-interruption blocks. The remaining 18 included an interrupting task during the memory delay. Interruption blocks were further subdivided depending on the temporal predictability of the interrupting task, having a fixed onset in 9 blocks (3 blocks each of 600 ms, 1400 ms, or 2400 ms after retro-cue onset, respectively), and a variable onset in the other 9 blocks (pseudo-randomized to occur equiprobably at 600 ms, 1400 ms, or 2400 ms after retro-cue onset). As such, the total number of trials where the interruption would appear at any of the three time points was equal between the corresponding fixed-onset blocks and across the variable-onset blocks.

The experiment was divided into three consecutive 30-minute sessions, with 5-minute breaks between each session. Unbeknownst to the participants, the sessions were further split into triplets, each containing one no-interruption block, one fixed interruption block, and one variable interruption block. The order of blocks within these triplets, as well as the order of triplets, was randomized. Before the start of each block, participants were informed whether they would have to respond to the interrupting task. However, they were not informed about the temporal predictability (i.e., fixed vs. variable) or about the three possible interference onsets (i.e., 600 ms, 1400 ms, 2400 ms). To familiarize themselves with the task, participants performed variable-onset interruption blocks until they reached an accuracy higher than 85% in the working-memory task and 90% in the interrupting task. Participants completed between two and five practice blocks to achieve this threshold.

### Apparatus and data acquisition

The experimental script was generated in Python using the PsychoPy package (version 2021.2.3; Peirce et al., 2019). Data acquisition took place in a dimly lit testing booth. Participants sat facing a monitor (27-inch Acer XB270H; resolution 1920×1080 pixels; refresh rate 100 Hz; screen width 60 cm) at a viewing distance of 100 cm. An eye tracker (EyeLink 1000, SR Research) was positioned on the table approximately 15 cm in front of the monitor. Before data acquisition and between blocks, eye position was calibrated using a 5-point calibration and validation method. Gaze was continuously tracked binocularly (or monocularly if binocular tracking was not possible) at a sampling rate of 1000 Hz, and participants were instructed to maintain central fixation throughout the trial.

EEG data were acquired using an SynAmps amplifier and Neuroscan data acquisition software (Compumedics Neuroscan, Abootsford, Australia). Sixty-one electrodes were positioned across the scalp using the international 10-10 positioning system. An additional electrode was placed on each mastoid (i.e., behind the left and right ear), and a ground electrode was positioned on the forehead. Vertical and horizontal electrooculography (EOG) were recorded using a bipolar montage, which involved placing electrodes beside the canthi of each eye as well as above and below the left eye. During acquisition, data were low-pass filtered at 500 Hz and digitized at a sampling rate of 1000 Hz. Electrode impedances were at least below 10 kΩ and ideally below 5 kΩ. Data were referenced to the left mastoid during data acquisition.

### Behavioral analysis

Behavioral data were analyzed in R Studio (Posit team, 2022), following the pipeline used in recent studies from our research group (Gresch, Boettcher, van Ede, et al., 2024; Gresch et al., 2021, 2022). During data pre-processing, trials were removed if reaction times (RTs; measured from probe onset to the first key press) exceeded 5000 ms or were 2.5 *SD* above the individual mean across the trials of all conditions. We additionally excluded trials in interruption blocks if participants did not respond, responded with the wrong key, or did not respond within 800 ms to the external task. Datasets with more than 15% of trials rejected during these pre-processing steps were removed from further analyses (*n* = 2). Moreover, we identified datasets with average reproduction errors (i.e., calculated by averaging the absolute difference between the angle of the target item and the reported angle) equal or higher than 45° in any of the conditions (*n* = 0). An additional two participants were excluded due to missing eye-tracking data. After these exclusion steps, datasets from 30 participants, with an average of 92.61% (*SD* = 2.67) of trials were retained.

The behavioral analysis examined the extent to which working-memory performance was impaired after completion of an external, interrupting task. We specifically asked whether working-memory representations were completely eradicated following interruption and, if not, whether these representations suffered compared to trials in which no interference occurred. Therefore, we tested whether reproduction errors differed from a chance level of 45° and compared reproduction errors between no-interruption and interruption blocks. We applied paired samples t-tests and report Cohen’s d as a measure of effect size.

To analyze the proportion of trials in which participants incorrectly reported the angle of the interrupters or made random guesses, we applied a probabilistic mixture model (Bays et al., 2009). The relative proportions of the non-target (i.e., interrupter) distribution and the guessing distribution were estimated for each participant. The mixture modelling analysis was performed using the mixtur package (version 1.2.1; Grange & Moore, 2022).

As in prior studies (Gresch et al., 2021; Lorenc et al., 2018; Mallett et al., 2020; Saito et al., 2023; Zhang & Lewis-Peacock, 2023), we quantified whether the reported working-memory orientation exhibited a bias either towards (i.e., attractive bias) or away from (i.e., repulsive bias) the angle of the interrupting items. To achieve this, we first corrected for general response biases by subtracting each participant’s mean reproduction error from their error on individual trials. Next, reproduction errors were sorted and binned using a moving-window approach (step size = 5°, bin width = 45°) based on the orientation of the target relative to the interrupter. The average response bias was calculated for each bin, resulting in a response-bias curve as a function of the target-interference angular difference. This approach provided response-bias values indicating the degree of deviation of the report from the target orientation, with negative values reflecting a clockwise shift and positive values reflecting an anticlockwise shift. Similarly, the target-interference angular difference was negative or positive depending on whether the interference was oriented clockwise or anticlockwise relative to the target. An attractive bias occurred when the response-bias sign matched the target-interference angular difference, while a repulsive bias was indicated when the signs were opposite. To statistically evaluate whether interrupters biased working-memory reports, we conducted a cluster-based permutation test with 1024 iterations, comparing the response bias curve against zero.

### Eye-tracking analysis

Pre-processing and analysis procedures followed previous work from our lab (Draschkow et al., 2022; Gresch, Boettcher, van Ede, et al., 2024; van Ede, Chekroud, & Nobre, 2019; van Ede et al., 2021). Eye-tracking data were first converted from edf to asc format, and subsequently read into R Studio (Posit team, 2022). Eye blinks were detected and interpolated ±100 ms around identified blinks. Data from the left and right eyes were averaged, yielding a single time course of horizontal gaze position (x-position) and vertical gaze position (y-position). Next, data were downsampled to 250 Hz (i.e., one sample every 4 ms), then epoched from -200 ms before to 4200 ms after the onset of the retro-cue. To focus on biases in fixational gaze behavior, trials were removed in which vertical or horizontal gaze positions exceeded ±2.75° (∼154 pixels) from fixation (i.e., further than half the distance to either memory item). We smoothed all time-course data over 15 samples (i.e., 60-ms smoothing window average) and we baselined each trial. We subtracted the mean value between -200 and 0 ms before the onset of the retro-cue from all time points. Eye-tracking data were recorded from 32 participants, with 30 included in the gaze analysis after excluding those with poor behavioral performance. On average, 94.11% (*SD* = 6.39) of trials were retained.

To compare lateral gaze biases across trials, we computed leftwards and rightwards gaze-position time courses within each onset condition relative to the retro-cue. To increase sensitivity and interpretability, we calculated the mean of the lateral gaze bias according to the location of the cued item (i.e., towardness). Towardness for each time step was given by the trial-average horizontal position in right-item trials minus the trial-average horizontal position in left-item trials (where position values left of fixation were negative) divided by two. A positive towardness indicates a gaze bias in the direction of the cued item. We used the towardness variable to determine the significance of gaze biases (compared to zero). We ran cluster-based permutation tests with 1024 permutations, effectively controlling for multiple comparisons across time points while maintaining high sensitivity.

For visualisation purposes, we constructed a heat map of gaze positions between 0 and 400 ms following interrupter onset, averaged across all onset conditions. For this analysis, we did not remove trials with gaze values exceeding more than half the distance of the placeholder centres. Two-dimensional kernel density estimations were obtained at a 150×150-pixel spacing on a 600×600-pixel square grid separately for cues indicating right and left memory items. Lastly, we subtracted density maps of left-from right-cued items which allowed us to illustrate the magnitude of the observed gaze bias.

### EEG pre-processing

EEG data were analyzed in Python using MNE-Python (version 1.4.2; Gramfort et al., 2013). The data were re-referenced to the common average, downsampled to 250 Hz, and filtered between 0.1 Hz and 50 Hz. Bad sensors were manually identified and interpolated. On average, 0.87 ± 0.82 (*M* ± *SD*) bad electrodes were identified and interpolated per participant. An independent component analysis (ICA) was used to identify and correct for artifacts associated with eye movements. For identifying ICA components associated with eye-movement artifacts, we correlated the time courses of all independent components with the measured horizontal and vertical EOG signal. Components with high correlation values and topographies characteristic of saccades or blinks were subsequently removed from the data. Data were then epoched from-200 to 4200 ms relative to the retro-cue onset and from -400 to 400 ms relative to the interruption response. No baseline correction was applied. We employed a generalized extreme studentized deviate (ESD) trial rejection approach to detect and exclude trials with high variance. In addition, we removed trials of interruption blocks in which participants did not respond to the interrupting task. On average, 29.07 ± 21.52 trials were removed per participant, but never more than 15% of all trials. Finally, we applied a surface Laplacian transform (Perrin et al., 1989) to increase the spatial resolution of spectral modulations under consideration (as also done in: Boettcher et al., 2021; Nasrawi et al., 2023; van Ede et al., 2019).

### Time-frequency analysis

We calculated time-frequency representations of power by convolving the data with Morlet wavelets across the frequency range of 3 to 40 Hz. For each frequency, we used a fixed 300-ms time window such that the number of cycles changed with each frequency.

To obtain a marker of visual selection, activity in predefined posterior visual electrodes (PO7/PO8) was contrasted between trials in which the memorized location of the cued item was contralateral versus ipsilateral to the electrode of interest. We expressed this contrast as a normalized difference (i.e., [(contra-ipsi)/(contra + ipsi)] × 100) and averaged it across the left and right electrodes. To derive time courses of lateralized visual activity, we averaged the contralateral versus ipsilateral response within the predefined alpha-frequency band (8-12 Hz). Topographies of lateralized visual activity were generated by contrasting trials in which cues indicated visual content in the right versus left hemifield, expressed as a normalized difference between right and left trials for each electrode (i.e., right minus left).

The analysis of motor selection mirrored the logic used for visual selection, but contrasted activity in predefined central motor electrodes (C3/C4) and between trials in which the prospective response hand was contralateral versus ipsilateral to the electrode of interest. Moreover, we computed time courses of motor literalization from the time-frequency contrasts by averaging across the predefined beta-frequency band (13-30 Hz). By independently manipulating item location and required response hand, we ensured that the EEG signatures of visual and motor selection were orthogonal in the trial average.

Statistical evaluation of EEG data used cluster-based non-parametric permutation, employing 1024 permutations and a cluster alpha of 0.05. This approach sidesteps the problem of multiple comparisons in the statistical analysis of EEG data (Maris & Oostenveld, 2007).

### Cross-correlations

We computed cross-correlations to test for potential latency differences between visual and motor time courses. Calculating cross-correlations involves shifting one time course relative to the other and computing the similarity (i.e., correlation) of the two signals at each shift (i.e., lag). The cross-correlation *z* at lag *k* is computed as:

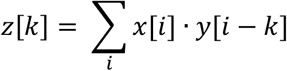

Each value of *z*[*k*] quantifies the similarity between the time series *x* and *y* when *y* is shifted by *k* indices. Here, *x*[*i*] represents the value of the array *x* at index *i*, while *y*[*i* − *k*] represents the value of the array *y* at index *i* − *k*, indicating that *y* has been shifted by *k*. Correlations are computed for lags *k* ranging from *k* = −*len*(*x*) + 1 (shifting *y* entirely to the right of *x*) to *k* = *len*(*y*) − 1 (shifting *y* entirely to the left of *x*). Hence, this range includes cases where *x* and *y* only partially overlap at the edges. The cross-correlation formula sums the product of corresponding values from one signal *x*[*i*] and the lagged version of the other signal *y*[*i* − *k*] over all indices *i*.

Since the raw cross-correlation values depend on the magnitude of the input signals, we normalized the results to make them independent of signal amplitude, ensuring the values are easier to interpret and statistically comparable. This normalization step scales the cross-correlation values to fall within the range [-1, 1]. Cross-correlations indicate how well the two signals match at each lag, that is, a high cross-correlation value at a specific time lag indicates a strong relationship between the two signals at that time lag.

To determine the lag at which the cross-correlation reached its maximum, we identified the individual peak in the cross-correlation for each participant. We then determined the lag corresponding to the maximum cross-correlation coefficient, which was subsequently tested against zero.

### Data and code availability

The experimental code, data, and analysis scripts are publicly available on the Open Science Framework (OSF): https://osf.io/c56eb/.

## Results

Participants performed a visual-motor working-memory task in which they memorized the orientation of two laterally presented colored bars (**Fig. 1A**). During the retention interval, a retro-cue indicated which of the two memory items would be relevant for an orientation-reproduction report at the end of the trial. In most blocks, following the retro-cue, working memory was interrupted by a perceptual discrimination task. Since this interrupting task could occur at three distinct onset times, we were able to determine whether reselection occurred immediately after responding to the interrupter or just in time when internal representations were required to guide behavior. Crucially, we also orthogonally manipulated the tilt (i.e., prospective response hand) and location of the cued item, allowing us to independently track sensory- and action-related working-memory content.

### Internal representations are disrupted but not lost

As our main interest in this experiment was to investigate the reselection of working-memory contents after task-relevant interruption, we first wished to confirm that participants were successful in reporting information from working memory following interruption. Our results clearly demonstrate that interruption did not eliminate working-memory representations. In both conditions, reproduction errors were very small (**Fig. 1B**; no-interruption trials: 10.451°± 0.220 [*M* ± *SEM*]; interruption trials: 11.885° ± 0.220) and significantly better than the chance level of 45° (no-interruption trials: *t*_(29)_ = 83.246, *p* < 0.001, *d* = 15.199; interruption trials: *t*_(29)_ = 73.595, *p* < 0.001, *d* = 13.436). This was further supported by the low number of trials in which participants used the incorrect response hand to indicate the working-memory orientation, with no significant difference observed between interrupter and no-interrupter trials (no-interruption trials: 2.231% ± 1.469; interruption trials: 1.670% ± 1.469; *t*_(29)_ = 1.478, *p* = 0.150, *d* = 0.270). Although working-memory performance was very precise in both conditions, reproduction errors were systematically larger when participants had to respond to the interrupter during the maintenance interval as compared to trials without external-task demands (**Fig. 1C**; *t*_(29)_ = 4.605, *p* < 0.001, *d* = 0.841). Thus, while the behavioral results indicate that interruption somewhat impaired working-memory performance, internal representations were not erased.

To examine how the interrupter task influenced the working-memory report, we analyzed reproduction biases. The results revealed that the internal representation of the probed orientation was systematically biased towards the orientation of the interrupting items (**Fig. 1D**; attractive bias; 1^st^ cluster *p* = 0.001, 2^nd^ cluster *p* = 0.001). Additionally, mixture modeling indicated that the proportion of guesses and swap errors with the interrupting items were very low (**Fig. 1E**; guesses: 0.028 ± 0.006; swaps: 0.009 ± 0.004), with the proportion of swaps being significantly smaller than that of random guesses (*t*_(29)_ = 3.533, *p* = 0.001, *d* = 0.645). Hence, the working-memory representation was inaccessible for recall in fewer than 3% of trials and replaced by the interrupters in fewer than 1% of trials. Taken together, participants were generally able to reselect the probed orientation, albeit with a slight bias towards the orientation of the interrupting stimuli.

### Internal selection during no-interruption trials

Before considering reselection of working-memory contents after interruption, we employed an EEG time-frequency analysis to investigate the dynamics of visual and motor selection after the retro-cue onset during no-interruption trials. While signatures of visual selection following retro-cues have been frequently observed (Gresch, Boettcher, Gohil, et al., 2024; Myers et al., 2015; Wallis et al., 2015), the question of whether motor selection similarly occurs during the prioritization of memorized information – but prior to the requirement to act on it – remains less explored (see also: Echeverria-Altuna et al., 2024; Ester & Weese, 2023; Schneider, Barth, Getzmann, et al., 2017). Establishing such an effect is an important proof of concept to ensure our design can detect the main effects of interest.

An established neural signature of visual selection is the lateralization of 8-12 Hz alpha-band activity in posterior electrodes contra-versus ipsilateral to the location of an attended item, both in perception (Gould et al., 2011; Gresch, Boettcher, Gohil, et al., 2024; Myers et al., 2015; Thut et al., 2006; Wallis et al., 2015; Worden et al., 2000) and working memory (Boettcher et al., 2021; Gresch, Boettcher, Gohil, et al., 2024; Myers et al., 2015; van Ede, 2018; Wallis et al., 2015). Similarly, the relative attenuation of beta-band activity in central electrodes contra-versus ipsilateral to the prospective response hand associated with a to-be-reported item has been linked with action selection (Boettcher et al., 2021; Ester & Weese, 2023; Nasrawi et al., 2023; van Ede, Chekroud, Stokes, et al., 2019; Zickerick, Kobald, et al., 2021).

In line with previous studies (Boettcher et al., 2021; Gresch, Boettcher, Gohil, et al., 2024; Myers et al., 2015; van Ede, 2018; Wallis et al., 2015), lateralization of neural activity relative to the visual location of the cued memory item showed a clear relative attenuation of alpha-band activity in contra-versus ipsilateral electrodes PO7/PO8 (**Fig. 2A**; cluster *p* < 0.001). The time course of the averaged 8-12 Hz alpha-band lateralization in the posterior electrodes revealed that the effect started early after retro-cue onset and was transient (Fig. 2C; cluster *p* < 0.001). Topographies show differences between trials in which the cued item occurred on the right versus left side of the display. They confirmed attenuation of alpha-band activity concentrated in the right and left posterior electrodes, consistent with visual selection (**Fig. 2E**).

**Figure 2.**
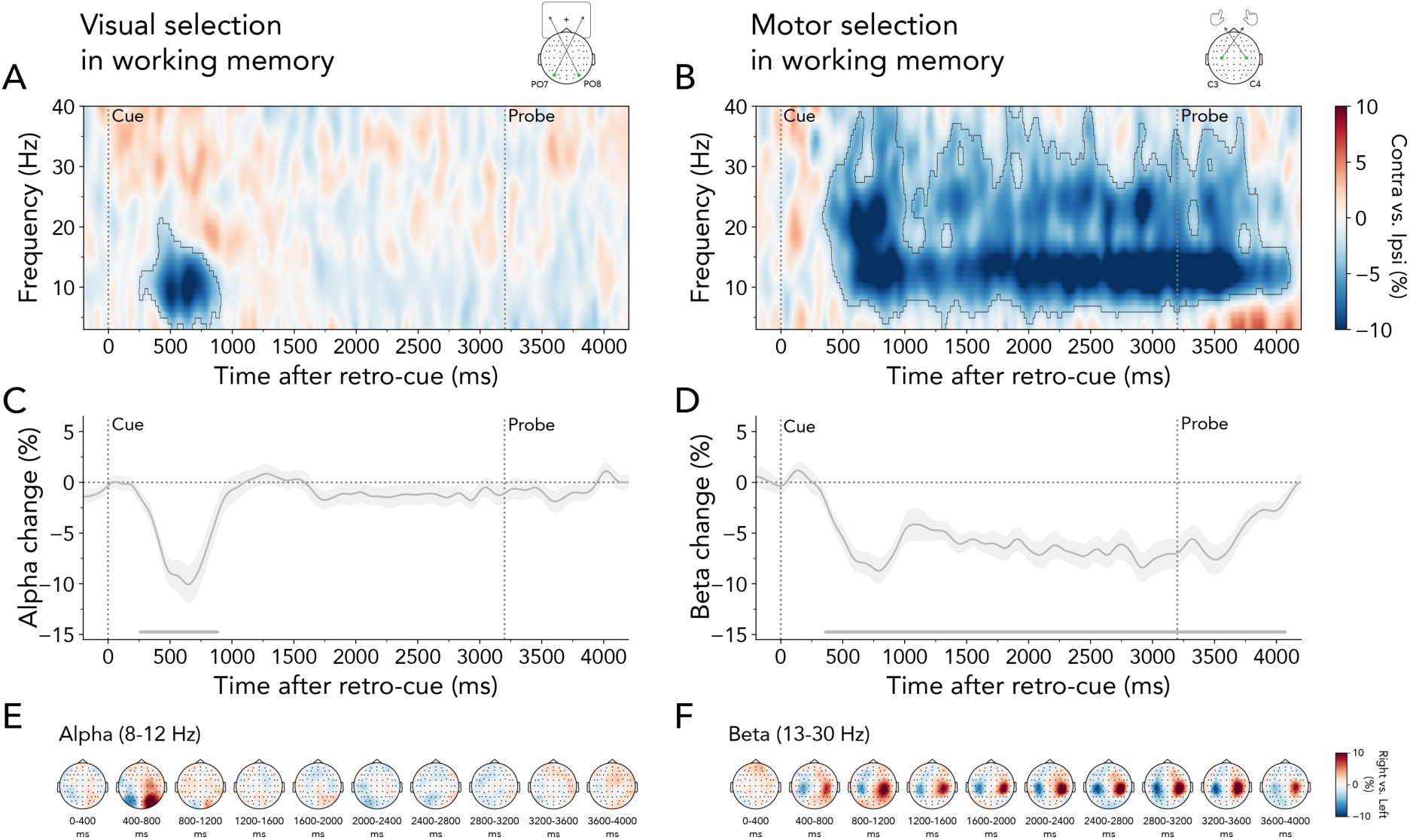
Visual and motor selection following retro-cue onset in no-interruption blocks. **(A)** Lateralized time-frequency representation contra-versus ipsilateral relative to the cued item’s location (left/right), calculated in electrodes PO7/PO8. The black outline indicates a significant cluster. **(B)** Same as (A) but relative to the response hand required to reproduce the cued item’s orientation (left/right), calculated in electrodes C3/C4. **(C)** Time course of lateralized visual activity at 8-12 Hz. The time course and shading show *M* ± *SEM* across participants. The horizontal line indicates a significant cluster. **(D)** Same as (C) but for motor activity at 13-30 Hz. **(E)** Topographies of the difference in 8-12 Hz alpha power between trials in which the retro-cue indicated the selection of the right or left item in working memory (right minus left), averaged in steps of 400 ms after retro-cue onset. **(F)** Same as (E) but showing the difference in 13-30 Hz beta power between trials in which the retro-cue indicated the right or left prospective response hand.

We also observed clear relative attenuation of beta-band activity (13-30 Hz) in electrodes C3/C4 contra-versus ipsilateral to the required response hand associated with the cued item (**Fig. 2B** and **D**; time-frequency representation cluster *p* < 0.001; time-course cluster *p* < 0.001). In contrast to the transient selection of visual information, action planning remained sustained following retro-cue onset, persisting until after probe presentation. Topographies confirmed that the beta-band modulations (13-30 Hz) between trials requiring a right-versus left-hand response were particularly prominent in the right and left central electrodes, consistent with action selection (**Fig. 2F**).

These data thus reveal that visual and response-hand-specific action selection occurred immediately following the retro-cue. While visual selection was transient, action-selection signatures remained sustained throughout the maintenance period. This pattern aligns with previous research (Echeverria-Altuna et al., 2024; Ester & Weese, 2023; Schneider, Barth, & Wascher, 2017), confirming that both visual and motor information can be prioritized promptly, even when memory-guided behavior is not immediately needed.

### Four possible scenarios of internal reselection following interruption

Having confirmed visual and motor selection following the retro-cue, we addressed our main questions: (1) *what* internal information is reselected following an interfering event which imposes additional task demands, and (2) *when* does this reselection occur? Key features of our design allowed us to disambiguate between several possible scenarios. Based on previous work, we considered two possible outcomes regarding the first question. One possibility is that after interruption, only motor plans need to be reselected (**Fig. 3, upper row**). For instance, during initial reselection, relevant feature values (e.g., the cued orientation) may be extracted, and corresponding motor plans programmed, with the cued location subsequently discarded. Consistent with this idea are recent findings, revealing dynamic transformations of internal representations into task-relevant codes that were shifted away from non-relevant attributes such as the spatial location of the memorized item (Henderson et al., 2022; Panichello & Buschman, 2021). Alternatively, if visual-spatial information is critical for the reselection of internal contents, both motor and visual information may be reinstated following interruption (**Fig. 3, lower row**).

**Figure 3.**
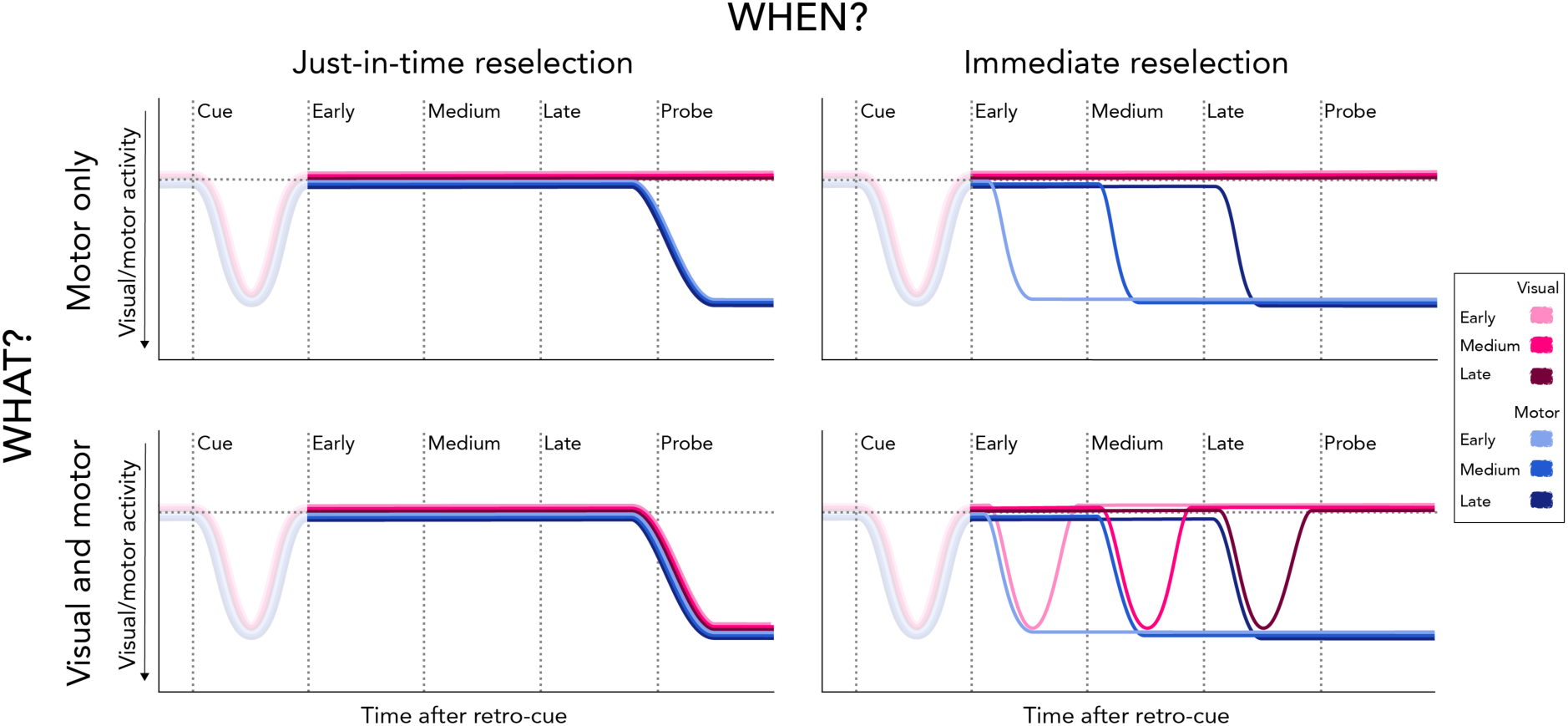
Four possible scenarios regarding the reselection of working-memory contents following task-relevant interference. **Top left:** Motor signatures could be reselected just in time when required to respond to the working-memory task. **Top right:** Motor signatures could become reselected immediately following the external, interrupting task. **Bottom left:** Visual and motor signatures could be reselected just in time when required to respond to the working-memory task. **Bottom right:** Visual and motor signatures could be reselected immediately following the external task. The magnitude differences in this figure (i.e., following the retro-cue) are intended solely to enhance the visibility of the different time courses.

An important feature of our design is the inclusion of three possible interrupter onsets. With these distinct onsets and a predictable probe time, we can determine whether information is reselected just before it is needed (i.e., around the expected time of the working-memory probe) or immediately after the interrupter. If memory reselection occurs just in time to guide performance, we would expect the shift towards internal representations to remain unaffected by the onset time of the external, interrupting task (**Fig. 3, left column**). However, if information is reinstated immediately after interruption, reselection onset should depend on the onset time of the interrupting task (**Fig. 3, right column**).

Considering the insights from no-interruption trials and prior research (Boettcher et al., 2021; Echeverria-Altuna et al., 2024; Ester & Weese, 2023; Gresch, Boettcher, Gohil, et al., 2024; Mok et al., 2016), we anticipated visual reselection to be transient, while motor reselection should remain more sustained across all scenarios. While various scenarios are also possible for the initial selection of visual and motor information prior to interruption, we primarily focus on the dynamics of reselection following interruption.

### Visual and motor reselection following interruptions

To address *what* internal information is reselected following interruption, and *when* this reselection occurs, we employed time-frequency analyses separately for each of the three interrupter onsets. We combined data from both the fixed- and variable-onset blocks, as there were no significant differences in lateralized visual and motor activity between the temporal-predictability conditions for early, medium, and late interruption onsets.

We observed relative alpha-band attenuation in electrodes PO7/PO8 contra-versus ipsilateral to the cued item’s memorized location for each of three onset conditions **(Fig. 4A**; early onset: 1^st^ cluster *p* < 0.001, medium onset: 1^st^ cluster *p* < 0.001, late onset: 1^st^ cluster *p* < 0.001; upper, middle, and lower panel, respectively). Initial alpha lateralization was succeeded by a second significant cluster of alpha-band attenuation after interruption (early onset: 2^nd^ cluster *p* = 0.004; medium onset: 2^nd^ cluster *p* = 0.025, 3^rd^ cluster *p* = 0.028; late onset: 2^nd^ cluster *p* = 0.009). These results were complemented by time courses of averaged 8-12-Hz alpha-band lateralization highlighting that significant attenuation occurred twice during the trial for all onset conditions (**Fig. 4C**). We can thus confirm that visual information of internal representations is reselected following task-relevant interference allowing us to rule out a purely motor-based account of memory reselection (i.e., **Fig. 3, upper row**). While the initial significant cluster appeared shortly after the retro-cue in all onset conditions, the second cluster emerged in a graded manner: first in the early-onset, followed by the medium-onset, and finally in the late-onset condition (early onset: 1^st^ cluster *p* < 0.001, 2^nd^ cluster *p* = 0.008; medium onset: 1^st^ cluster *p* < 0.001, 2^nd^ cluster *p* = 0.004; late onset: 1^st^ cluster *p* < 0.001, 2^nd^ cluster *p* = 0.006). These results suggest that mnemonic content is immediately reselected following interruption.

**Figure 4.**
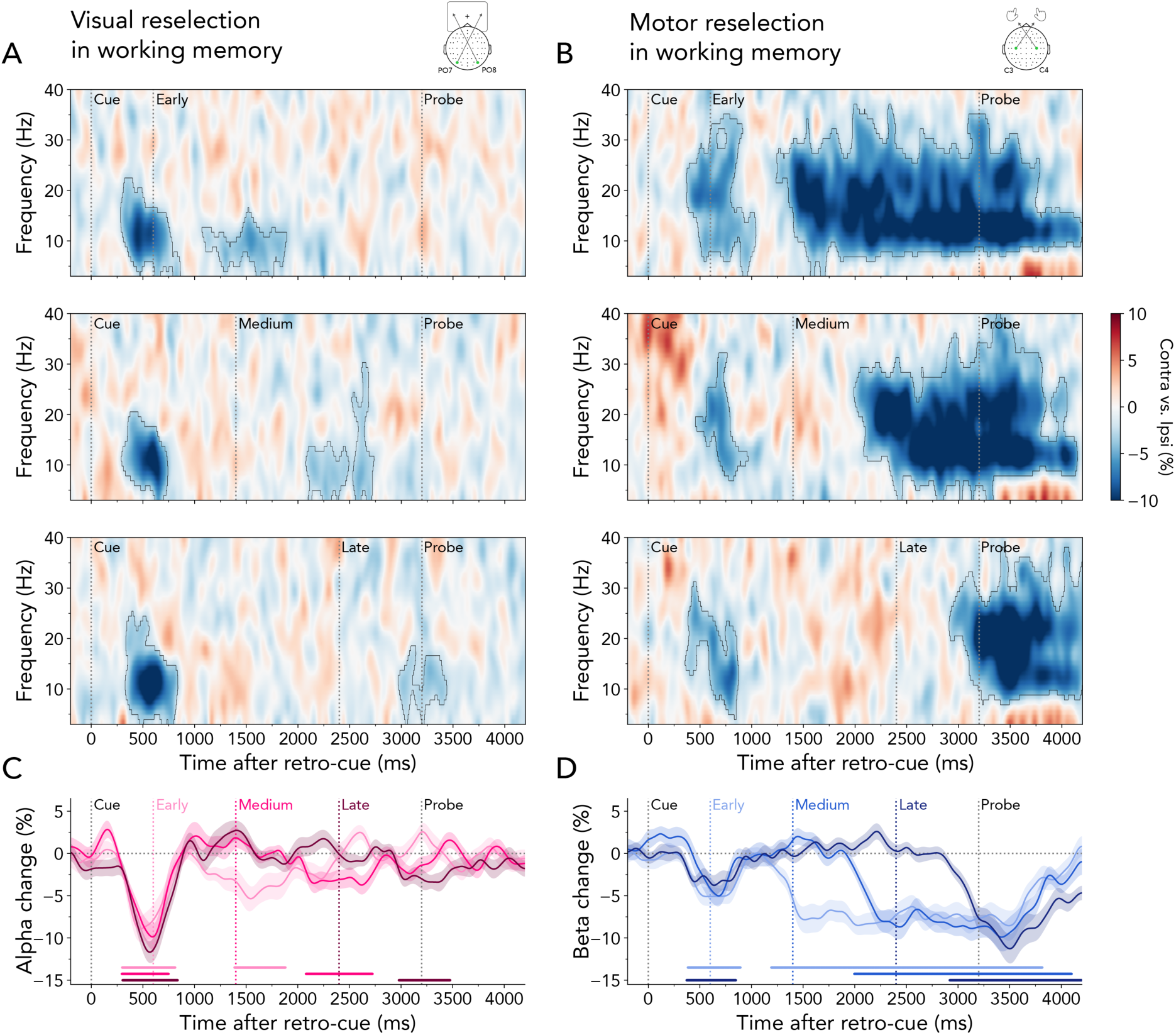
Visual and motor reselection following the interrupting task. **(A)** Lateralized time-frequency representation contra-versus ipsilateral relative to the cued item’s location (left/right), calculated in electrodes PO7/PO8. Top panel: early interrupter onset, middle panel: medium interrupter onset, bottom panel: late interrupter onset. Black outlines indicate significant clusters. **(B)** Same as (A) but relative to the response hand required to reproduce the cued item’s orientation (left/right), calculated in electrodes C3/C4. **(C)** Time courses of lateralized visual alpha-band activity (8-12Hz) for early, medium, and late interrupter-onset trials. Time courses and shadings show *M* ± *SEM* across participants. Colored horizontal lines indicate significant clusters. **(D)** Same as (C) but for motor beta-band activity (13-30 Hz).

Early motor signatures following the retro-cue showed a similar pattern as the initial visual selection. For all three interruption onsets, time-frequency representations revealed a relative attenuation of beta power in electrodes C3/C4 contra-versus ipsilateral to the response hand required to reproduce the tilt of the cued item (**Fig. 4B**; early onset: 1^st^ cluster *p* = 0.020, medium onset: 1^st^ cluster *p* < 0.001, late onset: 1^st^ cluster *p* = 0.010; upper, middle, and lower panel, respectively). The transient beta lateralization was followed by a more sustained lateralization, which varied temporally according to the onset of the interrupting task (early onset: 2^nd^ cluster *p* < 0.001; medium onset: 2^nd^ cluster *p* = 0.041, late onset: 2^nd^ cluster *p* < 0.001). The time courses of averaged lateralization in the 13-30 Hz beta-band confirmed a sequential pattern in the reselection of motor signatures, occurring earliest in early-onset trials and latest in late-onset trials (**Fig. 4D**; early onset: 1^st^ cluster *p* = 0.018, 2^nd^ cluster *p* < 0.001; medium onset: cluster *p* < 0.001; late onset: 1^st^ cluster *p* = 0.013, 2^nd^ cluster *p* < 0.001).

Together, these results yield two important insights. First, they provide compelling evidence for an initial selection of visual and motor information followed by immediate reselection after an external, interrupting task (i.e., **Fig. 3, bottom right**). Second, while the reselection of visual contents was short-lived, the reselection of motor information persisted for the remainder of the maintenance interval.

### Visual reselection biases gaze towards the location of the cued memory item

The primary aim of our study was to compare visual and motor reselection, with EEG data serving as the central metric for this investigation. Nonetheless, fixational gaze behavior can offer a rich source of complementary insights into the (re)selection of visuo-spatial contents (e.g., Gresch, Boettcher, van Ede, et al., 2024; van Ede, Chekroud, & Nobre, 2019). To this end, we analyzed lateral gaze biases. We combined right- and left-cued gaze time courses into a single ‘towardness’ metric for the three interrupter-onset conditions (**Fig. 5A**). Consistent with our EEG results, we observed spatial biases in gaze towards the retro-cued working-memory location across all three conditions (early onset: 1^st^ cluster *p* = 0.046; medium onset: 1^st^ cluster *p* = 0.047; late onset: 1^st^ cluster *p* = 0.024). Additionally, after interruption, gaze was promptly redirected towards the location of the cued memory item (early onset: 1^st^ cluster *p* < 0.001; medium onset: 1^st^ cluster *p* < 0.001; late onset: 1^st^ cluster *p* = 0.003). Density maps of gaze positions contrasting right- and left-cued trials show that this bias was primarily composed of small displacements towards the location of the cued item (**Fig. 5B**).

**Figure 5.**
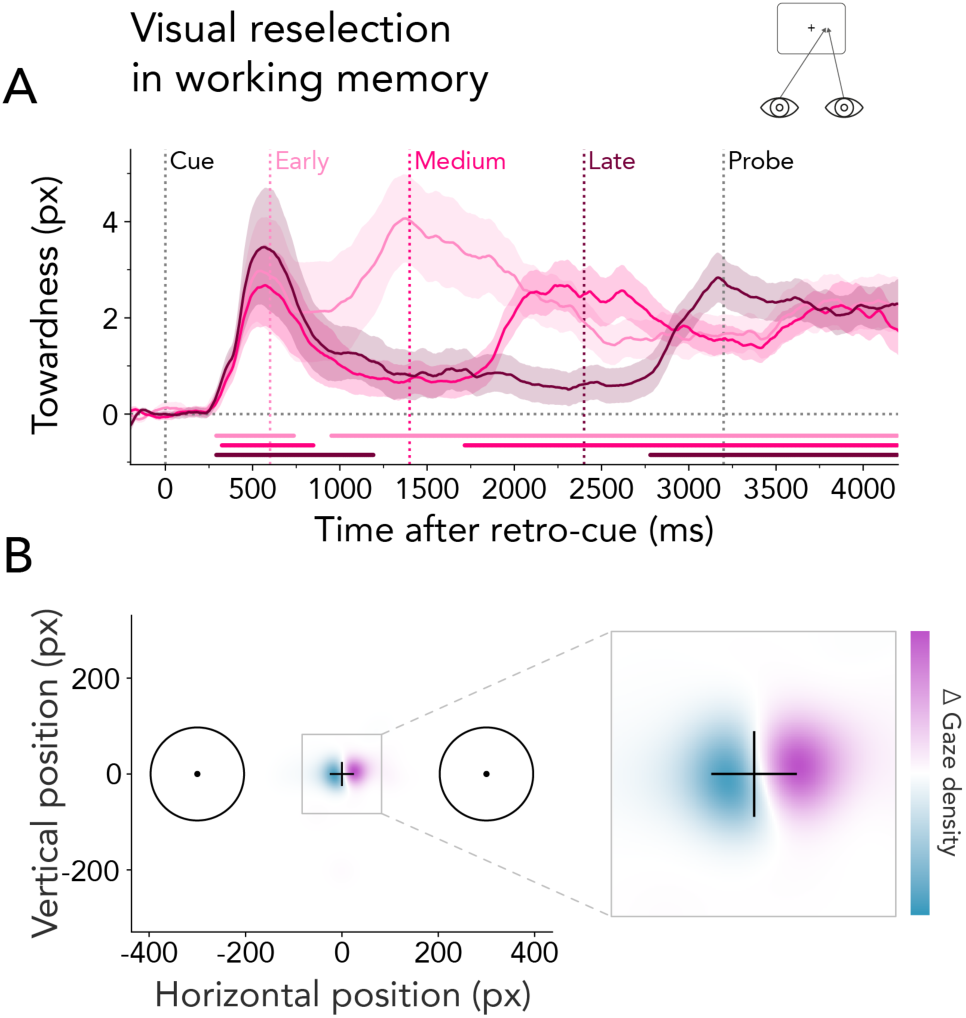
Visual reselection from working memory biases gaze towards the location of the cued item. **(A)** Average horizontal gaze bias expressed as towardness relative to the retro-cue onset for early, medium, and late interrupter-onset conditions. Time courses and shadings show *M* ± *SEM* across participants. Colored horizontal lines indicate significant clusters. **(B)** Difference in gaze density (Δ) following retro-cues indicating right versus left items (0–400 ms post interrupter across all conditions).

These results corroborate our EEG findings, providing additional evidence that participants initially selected visual information following the retro-cue and subsequently reselected this information after an interrupting task. Notably, a key difference emerged between the gaze bias and the reported alpha patterns. While relative alpha suppression was transient following early, medium, and late interrupters, the gaze bias was more sustained, persisting even after the probe. Recent studies have demonstrated that directional biases in fixational gaze behavior correlate with neural signatures of spatial attention but are not mandatory for neural modulation by spatial attention to occur (Liu et al., 2022). Our findings support this observation.

### Immediate internal reselection following interrupter responses

Our results highlight that the reselection of mnemonic information included both visual and motor signals and occurred immediately after the interrupting task, rather than just in time when working memory was needed to guide behavior. This prompts the question of how quickly participants shifted from the interrupter to the location and associated actions of internal contents. To address this, we combined all interruption trials and epoched the EEG data from-400 ms to +400 ms relative to the interrupter response. This epoching window was selected to avoid contamination of the results with the onset of the probe and the following working-memory response. The late interrupter appeared 2400 ms after retro-cue onset, with the probe following at 3200 ms. This provided an 800-ms window for participants to report the interrupter’s orientation. Given that RTs to the interrupting task in the late-onset condition were on average 401.542 ± 2.233 ms (*M* ± *SEM*), this window effectively isolates the neural activity linked to the reactivation of internal contents without interference from later events (i.e., the probe which appears at 3200 ms after retro-cue onset).

Immediately after the participants responded to the interrupter (∼20 ms after interrupter response), we observed a clear attenuation of alpha-band power in visual electrodes contra-versus ipsilateral to the location of the cued item (**Fig. 6A** and **C**; time-frequency representation cluster *p* < 0.001; time-course cluster *p* < 0.001). Analysis of beta-band lateralization relative to the required response hand yielded similar early signatures of motor reselection in central electrodes (**Fig. 6B** and **D**; ∼20 ms after interrupter response; time-frequency representation: cluster *p* < 0.001; time-course cluster *p* < 0.001). Visual and motor reselection had lateralized posterior and central topographies, respectively, within 0 to 400 ms after the interruption response (**Fig. 6C** and **D**).

**Figure 6.**
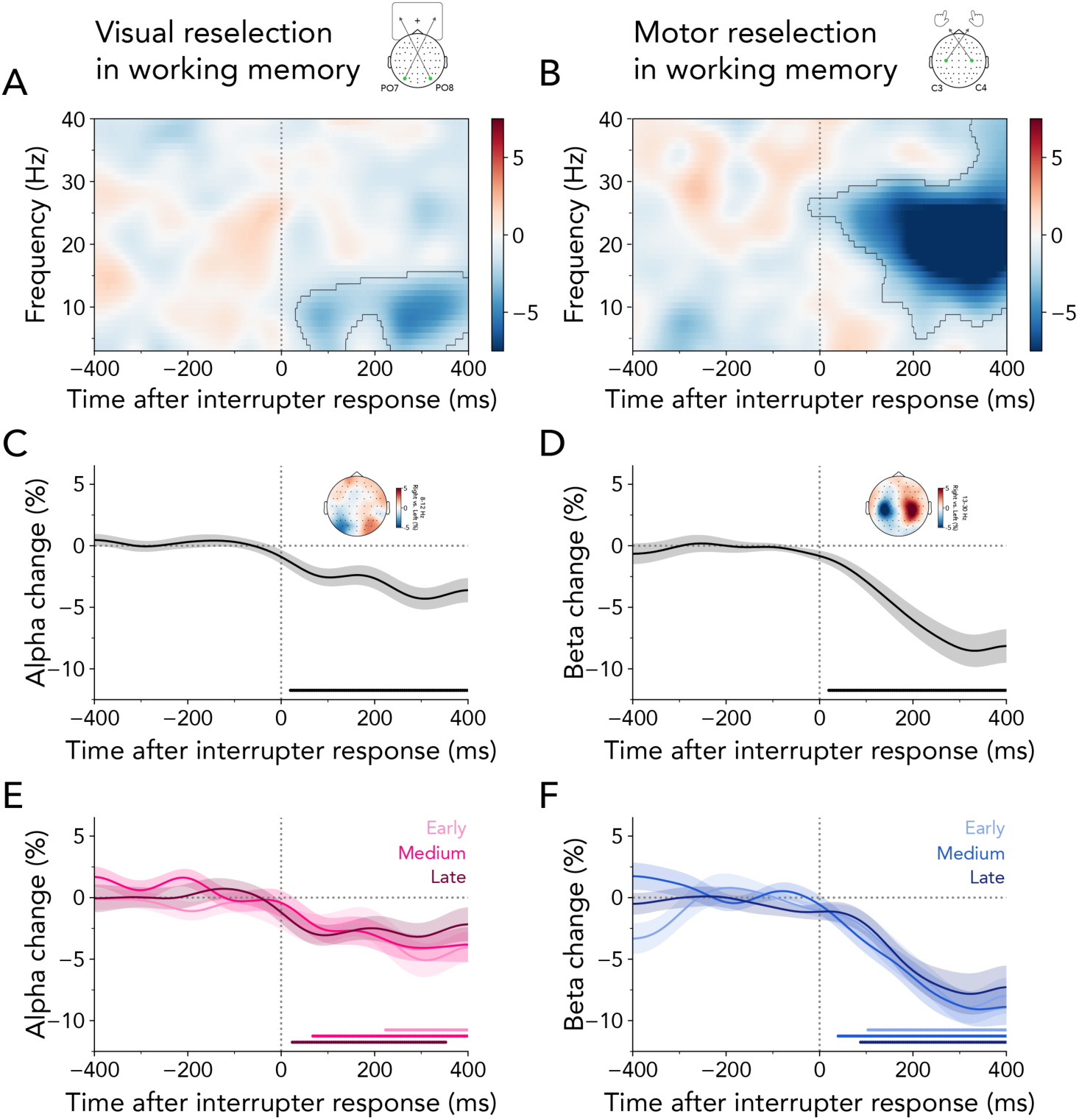
Visual and motor reselection occur immediately after the interrupter response. **(A)** Lateralized time-frequency representation contra-versus ipsilateral relative to the cued item’s location (left/right), calculated in electrodes PO7/PO8. The black outline indicates a significant cluster. **(B)** Same as (A) but relative to the response hand required to reproduce the cued item’s orientation (left/right), calculated in electrodes C3/C4. **(C)** Time course of lateralized visual activity at 8-12 Hz. Time course and shading show *M* ± *SEM* across participants. The black horizontal line indicates a significant cluster. The topography shows associated visual lateralization between 0 and 400 ms. **(D)** Same as (C) but for motor activity at 13-30 Hz. **(E)** Same as (C) but split by the three interrupter onsets. Colored horizontal lines indicate significant clusters. No significant differences were observed between the visual time courses. **(F)** Same as (D) but split by the three interrupter onsets. No significant differences were observed between the motor time courses.

To ensure that internal reselection was not predominantly driven by any specific onset condition, we divided trials according to the three interrupter onset times. We observed visual and motor reselection following the early (visual cluster *p* = 0.022; motor cluster *p* < 0.001), medium (visual cluster *p* = 0.002; motor cluster *p* < 0.001), and late interrupters (visual cluster *p*= 0.005; motor cluster *p* < 0.001), with no significant differences between these conditions (Fig. 6E and F). These findings collectively suggest that visual and motor reselection are tightly associated with the moment participants responded to the external task, further confirming that reselection is independent of the anticipated timing of memory-guided behavior.

### Visual and motor reselection overlap in time

Our results demonstrate that working-memory reselection engages both visual and motor brain areas immediately after responding to the interrupter. To examine the temporal relationship between visual and motor reselection with greater detail, we normalized visual and motor lateralization to their respective peak values. This analysis confirmed highly overlapping temporal profiles of visual and motor reselection (**Fig. 7A**).

**Figure 7.**
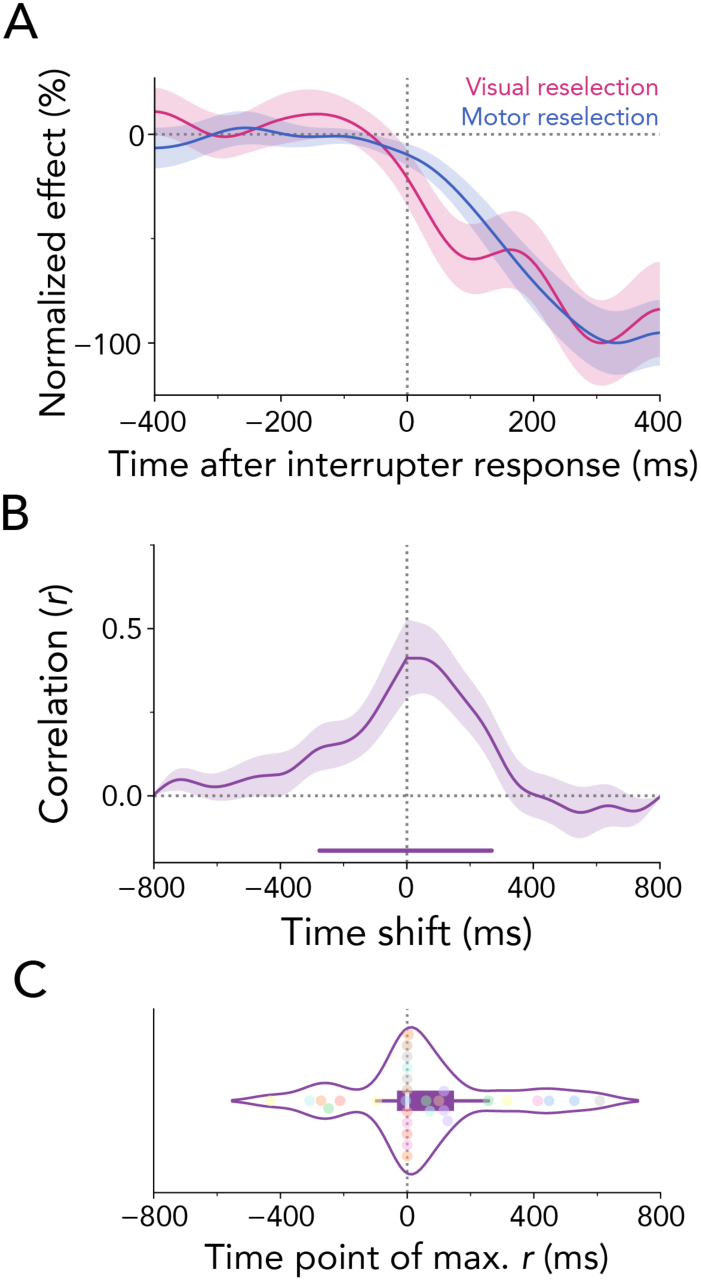
Parallel reselection of visual and motor attributes from working memory. **(A)** Normalized time courses for visual and motor selection. The visual reselection time course was obtained by averaging the item-location lateralization in selected visual electrodes between 8-12 Hz, while the motor reselection time course was obtained by averaging the response-hand lateralization in selected motor electrodes between 13–30 Hz. To increase visibility, the group-average time courses were normalized as a percentage of their peak value. Time courses and shadings show *M* ± *SEM* across participants. **(B)** Averaged cross-correlation coefficients between the time courses of visual and motor reselection. **(C)** Average lag at which the correlation coefficient reached its maximum. Dots represent individual participants.

One possible interpretation of this temporal correspondence at the group-average level is that it may reflect an artifact of averaging across participants, with some participants primarily showing visual reselection and others showing motor reselection signatures. To address this possibility, we performed a complementary analysis of temporal correspondence, calculating cross-correlations between the visual and motor reselection time courses for each participant. Cluster-based permutation tests revealed significant maximal correlations centered around the zero lag (**Fig. 7B**; cluster *p* < 0.001). Additionally, **Figure 7C** illustrates the time lags of maximal cross-correlations between visual and motor reselection time courses for individual participants. Importantly, no temporal shift was observed between visual and motor reselection relative to the zero lag (*t*_(29)_ = 1.225, *p* = 0.230, *d* = 0.224), consistent with the interpretation of a concurrent reselection of visual and motor attributes.

## Discussion

During natural behavior, we must often maintain internal representations in working memory while concurrently engaging with perceptual events in the external world. This raises a key question regarding how we reestablish internal focus following task-relevant interference. To this end, we investigated whether and when visual-spatial and motor attributes in working memory become reselected after engaging with an interrupting task. Our results provide evidence for the efficient and adaptable nature of memory-guided behavior. Specifically, visual and motor signatures reoccurred in tandem immediately following the interrupter response.

### What: Reselection of both visual and motor contents

Our study is the first to demonstrate that both visual and motor attributes are reselected following task-relevant interference. Specifically, our results revealed a transient lateralization in 8-12 Hz alpha power over posterior electrodes contra-versus ipsilateral to the cued item’s location, along with a sustained lateralization in 13-30 Hz beta power over central electrodes contra-versus ipsilateral to the prospective response hand following interruption. These findings align with commonly observed patterns of transient visual and sustained motor signatures during encoding, selection, and output gating (Boettcher et al., 2021; Echeverria-Altuna et al., 2024; Ester & Weese, 2023; Gresch, Boettcher, Gohil, et al., 2024; Mok et al., 2016; Myers et al., 2015; Wallis et al., 2015).

Motor reselection was expected since motor attributes were critical for memory-guided behavior in our task. In contrast, visual-spatial reselection is intriguing since it occurred despite spatial information being non-essential for working-memory performance. One might have expected the retro-cue to briefly direct visual attention to the hemifield where the cued item was previously presented, with subsequent reselection processes focusing solely on the motor and detailed sensory information (e.g., the item’s orientation), without additional reselection of the spatial location. Such a scenario would have been in line with evidence from electrophysiological and neuroimaging studies, demonstrating that neural signatures of internal representations are shifted into a task-oriented code that abstracts away from non-relevant attributes, such as spatial location (Henderson et al., 2022; Panichello & Buschman, 2021)

In contrast, we found that the previously cued spatial location is revisited, even when spatial information is not task-relevant and, in our case, has become entirely redundant following the initial item selection. These findings indicate that the spatial dimension may play a crucial role in scaffolding internal representations. Specifically, our results suggest that the sensorimotor mapping between the orientation and hand required to respond to the interrupting task disrupted the previously established response-ready representation necessary for memory-guided behavior. The reselection of the cued location after interruption may have been essential to reestablish the working-memory orientation-response link. Previous work has highlighted the special role space holds in binding sensory features (e.g., color and orientation) in memory (de Vries et al., 2023; Pertzov & Husain, 2014; Schneegans & Bays, 2017; Treisman & Zhang, 2006). Our results could suggest that location may also be important for binding response-related representations. Determining whether space serves as a necessary or sufficient condition for reselection processes is an intriguing avenue for future research. Moreover, it would be valuable to investigate the reselection of task-relevant visual features, such as orientation, in a task where the item’s orientation is not directly linked to the required action. This approach could help disentangle how spatial, motor, and feature attributes interact and unfold during reselection.

### When: Immediate reselection of visual and motor contents

The second goal of this study was to determine when internal attributes are reselected after interruption. Previous research has indicated an immediate return of alpha lateralization following the completion of an interrupting task embedded within a working-memory delay (Zickerick, Rösner, et al., 2021). However, the interrupting and working-memory tasks in this previous study were fundamentally different from one another. The working-memory task required participants to memorize and reproduce a tilted bar, whereas the interrupting task involved either a numerical or arithmetic comparison. Hence, transitioning between the interrupting task and the working-memory task also involved shifting from verbal to visual content. Given that there is less competition between verbal and visual contents in working memory (Baddeley, 1992; Brooks, 1968; Logie et al., 1990), it is reasonable to assume that a shift from verbal to visual contents may encounter less disruption than shifting between visual contents. In our study, we utilized a task design in which both working-memory and interrupting task engaged visual orientation judgments at the same peripheral locations, thus sharing similar stimulus-response mappings. Additionally, we systematically varied the onset time of the interrupting task which allowed us to pinpoint when reselection occurred. Our results revealed that for each of the three onset conditions, transient visual and sustained motor reselection occurred immediately after the completion of the interrupting task, rather than just in time before the probe onset. Moreover, our data suggest the concurrent availability of both visual and motor memory attributes following the interrupting response. Similarly, recent work has shown that when memorized visual information is probed to guide behavior, action plans can be selected from memory simultaneously with visual representations (van Ede, Chekroud, Stokes, et al., 2019). However, unlike the neural signatures observed in that study, our reselection results were not driven by an explicit cue prompting the retrieval of relevant mnemonic contents but rather reflect a natural endogenous reselection following an interrupting task.

The immediate reselection of visual and motor attributes may serve multiple purposes: First, it ensures that memories are prepared for use and remain readily accessible whenever internal content becomes relevant for guiding behavior. This may be a more computationally efficient process for the brain compared to engaging temporal expectations to reactivate internal content just in time. A second purpose of immediate reselection may be to enhance the robustness of memories, making them more resilient to subsequent decay.

Our task also allowed us to capture the natural flow of shifting attention from external to internal content without the use of explicit cues. Shifts to memory content in the current task were self-initiated, suggesting that the completion of an action can trigger the reselection of internal contents. However, a strictly serial approach to performing external tasks and reselecting internal information may not always be required or possible. Internal content may be reactivated while concurrently engaging with a secondary task, depending on the degree of competition for sensory and motor processing between the tasks and/or the time pressure to resume the working-memory task. For example, Zickerick, Rösner, et al. (2021) showed that demanding arithmetic interruptions can trigger even overlapping reselection of internal visual contents and interruption processing. Likewise, during sequential working-memory tasks, attention has been shown to preemptively shift to the next relevant internal content, even as actions guided by the currently relevant internal content are still ongoing (van Ede et al., 2021). The boundary conditions for determining when the processing of interrupting versus internal contents occurs serially or in parallel are an exciting avenue for further investigation.

Lastly, we should mention the possibility that visual and motor reselection may occur at different time points across individual trials. Latency variability at the single-trial level may be obscured by averaged signatures. Nonetheless, importantly, our results attest to there being no systematic differences between the timing of visual and motor reselection following interruption.

### Sustained or transient selection of motor contents?

Our results in the no-interruption condition replicated previously reported dissociations, demonstrating transient alpha suppression alongside sustained beta suppression following retro-cues (Echeverria-Altuna et al., 2024; Ester & Weese, 2023). Interestingly, visual inspection suggested that motor selection in interruption trials appeared more transient compared to no-interruption trials. The transient motor selection following the retro-cue, coupled with sustained reselection after the interrupter response, may indicate a biphasic profile of motor signatures. This aligns with previous findings on the encoding and subsequent preparation of prospective actions in working memory (Boettcher et al., 2021; Nasrawi et al., 2023). Moreover, it is striking that in interruption trials, initial action (and visual) selection was transient, irrespective of the interrupting task’s onset time. Safeguarding motor signatures by rapidly transitioning them into a latent state may be more efficient for memory-guided behavior than deactivating them shortly before the onset of an external task, which could risk disrupting internal representations. Taken together, our results suggest a level of context-dependent flexibility within internal representations, highlighting their inherently dynamic and adaptive nature.

### Conclusion

In this study, we investigated the temporal dynamics of reselecting working-memory contents following interruption, capturing the natural flow of shifting from sensory information to mnemonic representations in a task demanding both external and internal attention. Taken together, our findings revealed (1) that reselection occurs for both visual representations and their associated actions, (2) that reselection is time-locked to the interrupter response, irrespective of the anticipated timing of memory-guided behavior, and (3) that visual and motor reselection begin simultaneously but diverge in their dynamics, with visual reselection being transient while action planning is sustained.

## CRediT author contribution statement

Daniela Gresch: Conceptualization, Methodology, Software, Investigation, Formal analysis, Visualization, Writing – original draft. Larissa Behnke: Conceptualization, Methodology, Software, Investigation, Writing – review & editing. Freek van Ede: Writing – review & editing, Funding acquisition, Supervision. Anna C. Nobre: Conceptualization, Methodology, Writing – review & editing, Funding acquisition, Supervision. Sage E.P. Boettcher: Conceptualization, Methodology, Writing – review & editing, Funding acquisition, Supervision.

## Acknowledgements

This research was funded by a Wellcome Trust Senior Investigator Award (104571/Z/14/Z) and a James S. McDonnell Foundation Understanding Human Cognition Collaborative Award (220020448) awarded to A.C.N.; an Experimental Psychology Society Postdoctoral Fellowship awarded to S.E.P.B.; and an ERC Starting Grant from the European Research Council (MEMTICIPATION, 850636) and NWO Vidi Grant from the Dutch Research Council (14721) awarded to F.v.E. The research was also supported by the NIHR Oxford Health Biomedical Research Centre. The Wellcome Centre for Integrative Neuroimaging is supported by core funding from the Wellcome Trust (203139/Z/16/Z). For the purpose of open access, the authors have applied a CC-BY public copyright licence to any Author Accepted Manuscript version arising from this submission.

